# Elucidating the molecular interplay between LRRK2 and Rab GTPases

**DOI:** 10.64898/2026.04.30.722110

**Authors:** Hanwen Zhu, Liam Eade, Dario R. Alessi, Ji Sun

## Abstract

Gain-of-function mutations in *LRRK2* are a major cause of inherited Parkinson’s disease. *LRRK2* encodes a multidomain kinase, whose bidirectional interplay with Rab GTPases regulates critical cellular processes like lysosomal homeostasis. Certain Rabs, including Rab12 and Rab29, recruit LRRK2 to organelle membranes and stimulate its kinase activity; activated LRRK2 phosphorylates a subset of Rabs in their Switch-II motifs. Molecular basis governing selective Rab recognition by LRRK2 remains unclear. Here we structurally characterize LRRK2 interactions with representative Rab GTPases and identify three novel Rab-binding sites: site 4 for Rab8A/10, site 5 for Rab43, and site 6 for Rab5A, defining a total of six distinct binding sites that account for known LRRK2-interacting Rabs. Additionally, we elucidated the binding site of GABARAP, an ATG8 member that recruits LRRK2 to stressed lysosomes. Our findings provide a framework for therapeutic targeting of LRRK2 recruitment for Parkinson’s.

## INTRODUCTION

Parkinson’s disease (PD) is the second most common neurodegenerative disorder, affecting 1–2% of elderly population worldwide^1^. Missense mutations in the gene encoding leucine-rich repeat kinase 2 (LRRK2) are among the most common genetic predisposition for late-onset PD, accounting for ∼5% of familial and ∼1% of sporadic cases^2–4^. Over 250 LRRK2 variants have been identified, with most pathogenic mutations showing enhanced kinase activity^5–7^. Moreover, elevated activity of wild-type LRRK2 has also been linked to a higher risk of idiopathic PD, which comprises ∼90% of all PD cases^8^. Given that LRRK2 hyperactivity is implicated in both genetic and idiopathic PD, pharmacological inhibition of LRRK2 kinase activity has emerged as a promising therapeutic strategy for PD treatment^9–11^.

LRRK2 is a 286-kDa multidomain protein, including ARM, ANK, LRR, ROC, COR, KIN and WD40 domains, and has both kinase and GTPase catalytic activities^12^. LRRK2 regulates various cellular pathways implicated in PD, such as ciliogenesis, vesicle trafficking, autophagy, lysosomal function, and mitochondrial homeostasis^12–15^. Dysregulation of LRRK2 perturbs these processes and increases PD risk. Recent structural and functional studies have revealed the mechanisms governing LRRK2 self-assembly, autoinhibition, membrane recruitment and activation, filamentation on microtubules, and pharmacological inhibition by kinase inhibitors^16–23^, providing detailed mechanistic insights into its regulation at near-atomic resolution. Nevertheless, the precise molecular basis by which pathogenic LRRK2 mutations drive PD remains elusive.

The interplay between LRRK2 and Rab GTPases is of central importance and is widely regarded as a critical driver of pathogenesis in LRRK2-associated PD^14, 24–26^. Rab GTPases are master regulators of intracellular vesicle transport, membrane fusion and endolysosomal biology, and their dysfunction is tightly linked to PD pathology^24, 27^. Rab GTPases function both upstream and downstream of LRRK2^28, 29^, within a “Rab-LRRK2-Rab” signaling cascade. Members of the Rab32 subfamily, including Rab29, Rab32 and Rab38, can recruit and activate LRRK2 at specific organelle membranes^30–39^; while under stress, Rab12 recruits LRRK2 to damaged lysosomes and stimulates its kinase activity^40, 41^. We and others have shown that Rab29/32 recruit LRRK2 through its ARM9-ARM10 region (site 1)^17^, whereas Rab12 engages with the ARM6-ARM7 region (site 3)^19, 41^. LRRK2 recruited to these membranes becomes activated through mechanisms that remain incompletely understood and subsequently phosphorylates multiple Rab GTPases, including Rab3A/B/C/D, Rab8A/B, Rab10, Rab12, Rab29, Rab35, and Rab43, *in vitro* and/or *in vivo*^25, 26^. It should be noted that although LRRK2 efficiently phosphorylates Rab5A/B/C at the Switch-II motif in vitro^25^; to date there is no compelling evidence that Rab5 isoforms are phosphorylated at this site *in vivo* in a LRRK2-dependent manner. Phosphorylated Rab8A (pRab8A) and Rab10 (pRab10) are widely used as robust biomarkers for LRRK2 kinase activity. Additionally, pRab8 can bind the N-terminal region of LRRK2 (site 2), establishing a positive feedback loop^42^.

Disruption of the “Rab-LRRK2-Rab” cascade could influence key cellular pathways, including ciliogenesis, autophagy, endolysosomal system, and phagosome maturation^14, 43–48^. For example, engagement of LRRK2 by Rab12 or the presence of pathogenic LRRK2 mutations promotes phosphorylation of Rab10 and Rab8A thereby suppressing ciliogensis^25, 49^. Elucidating the molecular basis underlying LRRK2-Rab is therefore essential for understanding LRRK2 function and its contribution to PD. A central unresolved question is how LRRK2 selectively phosphorylates only a subset of Rab GTPases and engages Rab32/38, despite the human genome encoding approximately 70 Rabs—more than 40 of which harbor a Switch-II residue that could, in principle, serve as an LRRK2 phosphorylation site^24, 50^.

Here we analyzed the molecular interplay between LRRK2 and Rab GTPases (14 substrates plus Rab32 and Rab38), combining cryo-electron microscopy (cryo-EM), computational modeling, and biochemical assays. We determined high-resolution cryo-EM structures of LRRK2 in complexes with substrates Rab8A, Rab10, Rab43, as well as Rab5A. These structures revealed three previously unidentified Rab-binding sites on LRRK2 (site 4-6). Site 4 spans the COR, KIN, and WD40 domains of LRRK2 and accommodates Rab8A or Rab10. This binding interface is compatible with both the monomeric and dimeric conformations of inactive LRRK2. However, in the active conformation, Rab10 can engage site 4 only in the monomeric form and is incompatible with the dimeric state. Site 5 is positioned within the ARM13–15 and ANK domains and binds Rab43, whereas site 6 involves ARM12–14 domain and engages Rab5A. We also structurally characterized the interaction between LRRK2 and GABARAP, an ATG8 family member that may provide parallel or cooperative platforms for LRRK2 recruitment and activation at stressed lysosomes. The GABARAP binding interface is close to site 2, consistent with prior biochemical characterization and modeling studies^51^. Collectively, these findings define the molecular determinants of LRRK2–Rab interactions and establish a structural framework for the rational therapeutic targeting of these interfaces.

## RESULTS

### Structure of Rab10–LRRK2 complexes and the Rab-binding site 4

We first structurally characterized the molecular interaction between LRRK2 and Rab10. By trapping the globular G-domain of Rab10 in a GTP–bound state (Q68L; residues 1–181), we resolved the cryo-EM structures of Rab10–LRRK2 complexes, following a similar approach as we used for the Rab12–LRRK2 complex^19^. The Rab10–LRRK2 complexes were captured in both the LRRK2 monomeric and dimeric states at nominal overall resolutions of 4.1 Å and 3.3 Å, respectively (**Extended Data Fig. 1a-d and Table S1**). For the LRRK2 dimeric state, *C*2 symmetry was imposed during data processing, and focused 3D refinement was performed to improve the cryo-EM density of the N-terminal region. Rab10 bound LRRK2 with a 2:1 stoichiometry in the monomeric Rab10–LRRK2 complex, and 4:2 stoichiometry in the dimeric complex (**Extended Data Fig. 1a**), different from the 1:1 stoichiometry seen for Rab12–LRRK2^19^. The LRRK2 conformation in the Rab10–LRRK2 complex is nearly identical to that of LRRK2-alone (**Extended Data Fig. 1e**) and within the Rab12–LRRK2 complex (**Extended Data Fig. 1f**)^16^, consistent with an overall inactive state. The GTP analog–bound Rab10 conformation resembles that of Rab29 and Rab12 in their respective LRRK2 complexes (**Extended Data Fig. 1g**)^17, 19^.

In the full-length inactive LRRK2 monomeric and dimeric conformation, Rab10 is bound to two distinct sites, namely site 3 and site 4 (**Fig. 1a**). Site 3 recapitulates the previously reported Rab12–LRRK2 interface^19^, with the Switch I–Interswitch–Switch II region of Rab10 making contacts with the ARM6–7 repeats of LRRK2 (**Extended Data Fig. 1h**). In the dimeric conformation, the novel site 4 is located primarily within the C-terminal half of LRRK2, with Rab10 nesting within a cavity formed between the dimer interface (**Fig. 1a**). Site 4 involves three interaction interfaces: (i) the COR-B subdomain and kinase N lobe of LRRK2 protomer A engages with the Switch I–Interswitch–Switch II motifs of Rab10 in a canonical Rab–effector interaction mode (**Fig. 1b, d**); (ii) the ANK domain of LRRK2 protomer B interacts with the CDR5 region of Rab10 (**Extended Data Fig. 3**, **Fig. 1b, e**); and (iii) the COR and WD40 domains of LRRK2 protomer B, together with the COR-B subdomain of LRRK2 protomer A interact with the CDR1, Switch I−Interswitch, and C-terminus of Rab10 (**Fig. 1b, c**). In this third interface, the first five residues of Rab10 sit at the cleft formed between COR-A and COR-B subdomains of two LRRK2 protomers (**Fig. 1b, c**). Together, site 4 interfaces bury approximately 1600 Å^2^ (**Fig. 1b**), substantially exceeding the buried surface areas observed in the Rab29/32–LRRK2 (∼800 Å^2^) and Rab12–LRRK2 (∼530 Å^2^) complexes.

**Fig. 1.**
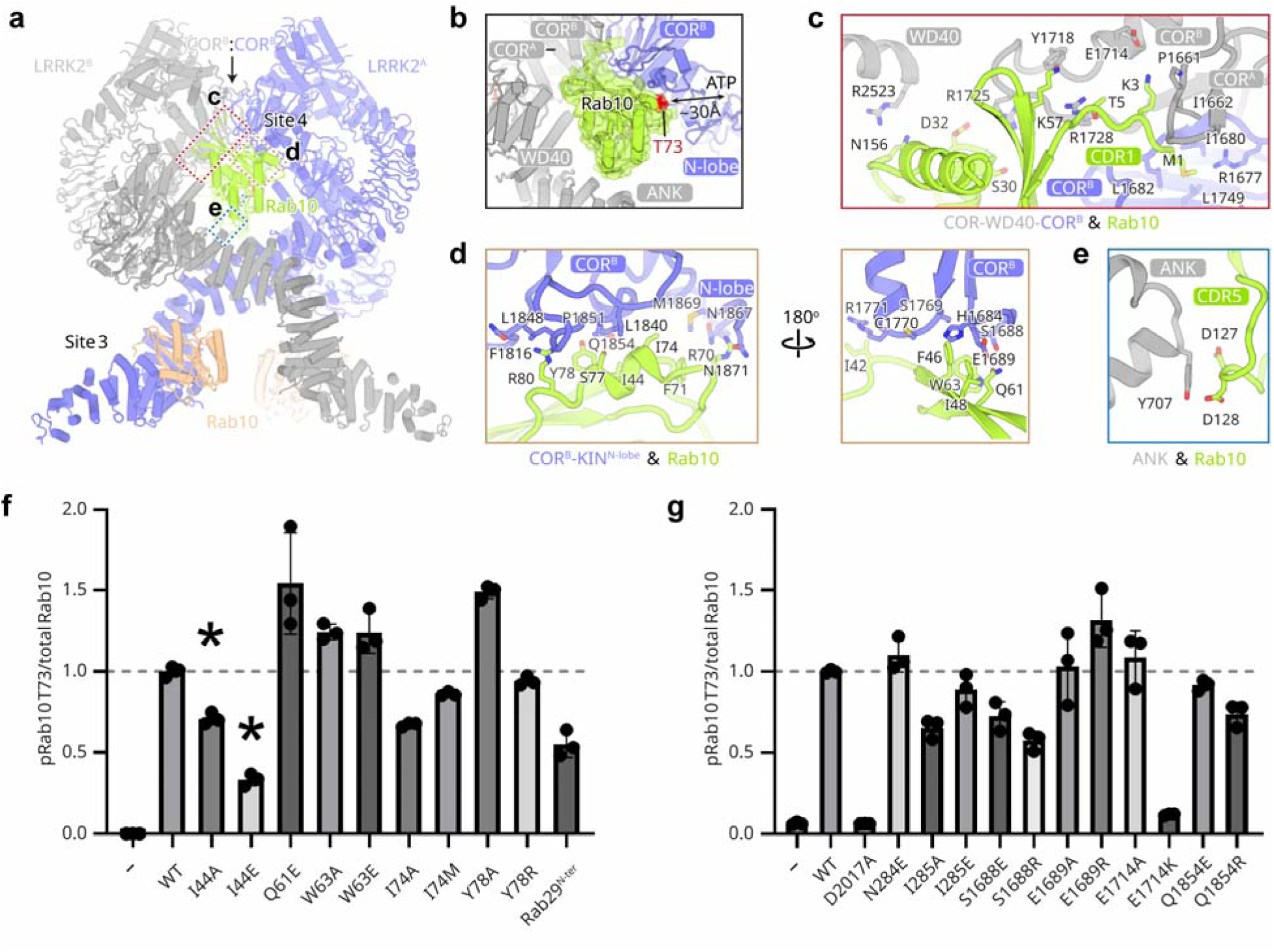
| Structural details and biological relevance of the Rab10–LRRK2 complex. **a**, Cryo-EM structure of the Rab10–LRRK2 complex in the LRRK2 dimer state. Color code: Rab10, limon and light orange; LRRK2, slate and grey. LRRK2 sites 3 and 4 engaged by Rab10 are indicated. **b,** Zoom-in view of the LRRK2–Rab10 interaction. Rab10-interacting domains in LRRK2 are labeled. The distance between the Rab10 Thr73 site and the LRRK2 active site is indicated. **c-e,** Interfaces between Rab10 and the LRRK2 COR/WD40 domains (**c**), the LRRK2 COR-B subdomain and kinase N-lobe (**d**), and the LRRK2 ANK domain (**e**). Side chains of interface residues are shown as sticks. **f-g,** Quantification of the immunoblotting data shown in Extended Data Fig. 2a, c. Data are presented as ratios of pRab10-Thr73/total Rab10, normalized to the average of LRRK2 WT values. The data shown are the mean ± SD of three determinations. * = impaired protein expression or stability.

In the Rab10–LRRK2 monomeric complex, Rab10 also binds LRRK2 at sites 3 and 4. However, at site 4, the interaction between CDR5 of Rab10 and ANK and WD40 domains of the neighboring LRRK2 subunit is absent. The interaction between Rab10’s CDR1 and LRRK2 may be partially retained, but the resolution in this region is insufficient for a definitive structural model.

To assess the functional significance of the Rab10–LRRK2 interface at site 4, we introduced interface mutations and evaluated their effects on cellular LRRK2 kinase activity by quantifying phosphorylation of Rab10-pThr73, Rab12-pSer106, LRRK2-pSer1292, and LRRK2-pSer935. Most mutations, whether on LRRK2 or Rab10, had little effect on Rab12-pSer106, LRRK2-pSer1292, or LRRK2-pSer935, with the notable exception of LRRK2 E1714K, which decreases Rab12-pSer106 level (**Extended Data Fig. 2b, d**). In contrast, Rab10-pThr73, was significantly reduced by three Rab10 mutants (**Fig. 1f; Extended Data Fig. 2a**): (i) I44A/E, which reduced pThr73 but also lowered total Rab10 levels, indicating stability impairment; (ii) I74A, located next to Thr73 and likely disrupting kinase-substrate recognition; and (iii) an N-terminal chimera in which the first five residues of Rab10 were replaced with the Rab29’s N-terminal loop, a region not implicated in LRRK2 binding^17^ (**Fig. 1f; Extended Data Fig. 3**). Sequence alignment revealed that Lys3 and Thr5 within this N-terminal motif are conserved only in Rab10 and Rab8A/B but not in Rab29 or other Rabs, suggesting this short N-terminal sequence may selectively anchors Rab10 and potentially Rab8A/B at site 4 (**Extended Data Fig. 3**). Corresponding LRRK2 mutations at Glu1714, forming a salt bridge with Rab10 Lys3 in the N-terminal loop (**Fig. 1c, g; Extended Data Fig. 2c**) or at Ser1688, contacting the Switch I–Interswitch–Switch II region of Rab10 (**Fig. 1d, g; Extended Data Fig. 2c**), reduced Rab10-pThr73 levels. Together these data suggest a functional importance of site 4 for efficient LRRK2-mediated phosphorylation of Rab10.

### Rab10 binding changes LRRK2 kinase conformation

Rab10 binding to the LRRK2 dimer interface triggers some global conformational rearrangement. When one protomer from LRRK2-alone structure is superimposed as a reference, the opposing protomer rotates outward by ∼15° to accommodate the intercalated Rab10 molecule (**Fig. 2a**). Within each protomer, Rab10 binding further induces a small clockwise rotation of the COR-B subdomain and the kinase N-lobe relative to the kinase C-lobe (**Fig. 2b**).

**Fig. 2.**
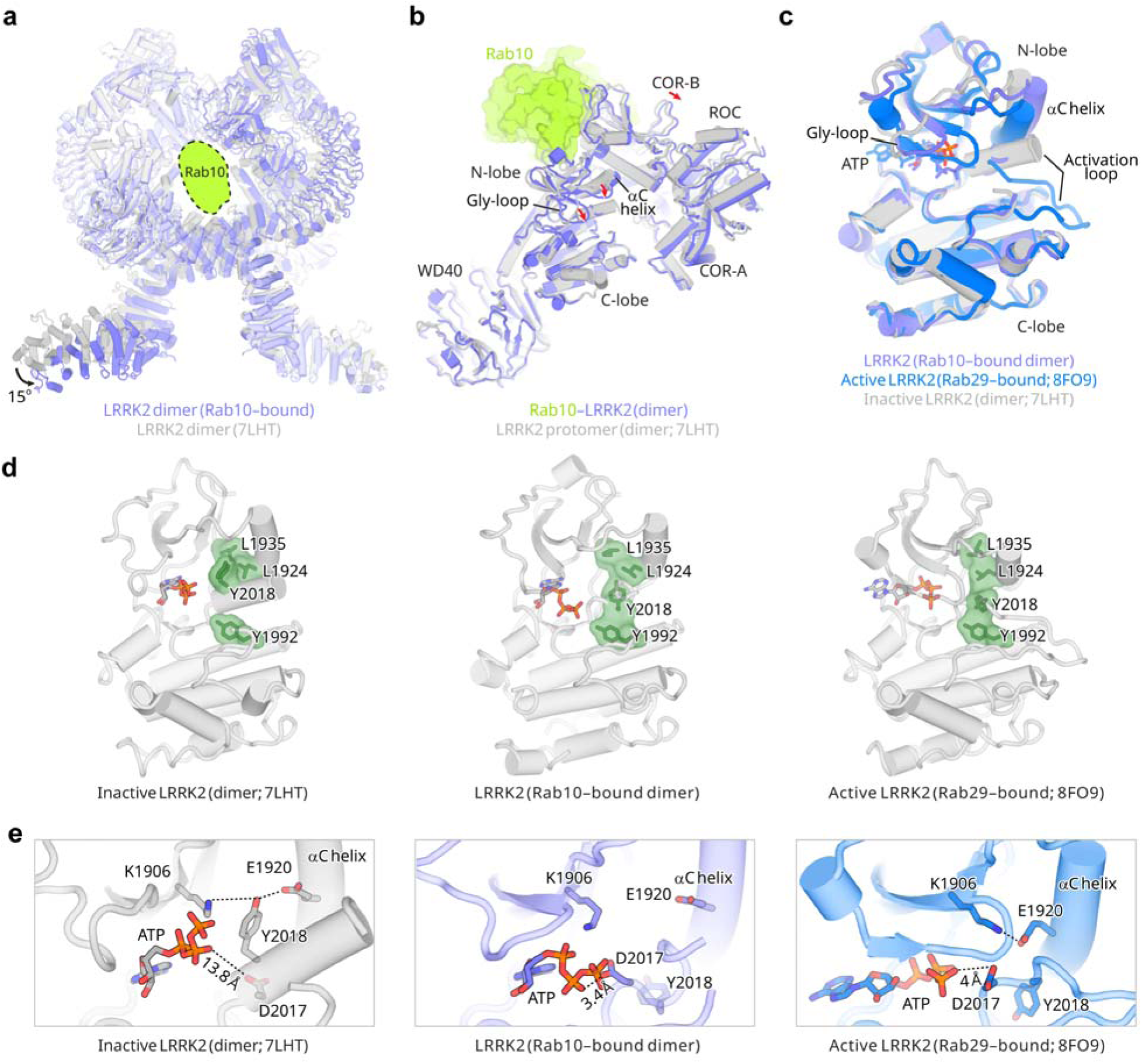
| The inactive intermediate kinase domain of Rab10-bound LRRK2. **a**, Structural comparison of the Rab10-bound LRRK2 dimer and the LRRK2 alone dimer (PDB 7LHT). The Rab10-bound LRRK2 dimer and LRRK2-alone dimer are colored in slate and grey, respectively. Rab10 binding at LRRK2 site 4 is indicated with a dashed oval. The overall conformational change of one LRRK2 protomer relative to the opposing LRRK2 protomer upon Rab10 binding is indicated with a red arrow. **b,** Conformational changes in the C-terminal halves of LRRK2 upon Rab10 binding. The movements of the LRRK2 COR-B subdomain and key structural elements in the LRRK2 KIN domain are indicated with red arrows. **c,** Superposition of the KIN domains of Rab10-bound LRRK2, active LRRK2 (PDB 8FO9), and inactive LRRK2 (PDB 7LHT). Key structural elements of the KIN domains are labeled. **d,** R-spine in the KIN domains of inactive LRRK2 (left; PDB 7LHT), Rab10-bound LRRK2 (middle), and active LRRK2 (right; PDB 8FO9). The four residues forming the R-spine (L1935, L1924, Y2018, and Y1992) are shown as green surfaces. **e,** Key catalytic residues in the KIN domains of inactive LRRK2 (left; PDB 7LHT), Rab10-bound LRRK2 (middle), and active LRRK2 (right; PDB 8FO9) with side chains shown as sticks. Distances between the side chain of D2017 and the phosphate group of the ATP are labeled.

This global rearrangement of LRRK2 is accompanied by localized conformational changes in the kinase domain, resulting in an intermediate inactive state. The αC helix and Gly-loop in the N-lobe shift toward the C-lobe, adopting an intermediate conformation that exhibits hybrid structural features between the inactive and active states (**Fig. 2c**). In this intermediate state, the regulatory spine (R-spine), a hallmark of the active kinase state, formed by Leu1935, Leu1924, Tyr2018, and Tyr1992, began to emerge (**Fig. 2d**). Notably, Tyr2018 within the “DYG” motif flipped inward into a “DYG-in” conformation resembling the active state, in contrast to the “DYG-out” conformation observed in the inactive kinase. Asp2017 formed a hydrogen bond with the ATP phosphate group, which is essential for catalysis (**Fig. 2e**). On the other hand, the Lys1906–Glu1920 salt bridge remained broken, and the activation loop was partially disordered, features consistent with an overall inactive state (**Fig. 2e**). Collectively, these structural observations indicate that Rab10 binding stabilizes an inactive intermediate conformation of LRRK2. In addition, Rab10 Thr73 is ∼30 Å away from the LRRK2 active site (**Fig. 1b**), a separation incompatible with phosphorylation in the captured structure; indicating that substantial additional conformational rearrangements would be required to enable Rab10 phosphorylation. Alternatively, site 4 could serve as a Rab-docking site, with phosphorylation occurring at a distinct, as yet unknown catalytic engagement site.

Site 4 is adjacent to a recently characterized secondary docking site for 14-3-3^52^. The scaffold protein 14-3-3 binds to phosphorylated LRRK2 at Ser910 and Ser935^53, 54^ and interacts secondarily with the COR-B subdomain, stabilizing an inactive LRRK2 conformation^52, 55^. Structural superposition analysis revealed potential steric clashes between Rab10 and 14-3-3 binding. In the monomeric Rab10–LRRK2 complex, the Rab10-binding interface at site 4 potentially overlaps with the 14-3-3 secondary interface on COR-B (**Extended Data Fig. 1i**); in the dimeric Rab10–LRRK2 complex, 14-3-3 would sterically clash with either the second LRRK2 protomer or the Rab10 bound at site 4 (**Extended Data Fig. 1j**). These observations suggest that Rab10 binding at LRRK2 site 4 could displace 14-3-3, potentially relieving 14-3-3–mediated inhibition and facilitating LRRK2 activation.

### Rab10 binding on site 4 is compatible with active monomeric LRRK2

We next examined whether Rab10 binding is compatible with the active conformation of LRRK2. We determined the cryo-EM structure of the Rab10 bound to LRRK2^RCKW^ in the presence of inhibitor LRRK2-IN-1 and DARPin E11 (**Extended Data Fig. 4 and Table S1**). Here, the LRRK2^RCKW^ construct lacks the N-terminal ARM-ANK-LRR domains that shield the kinase active site and become flexible upon activation^17, 18^; LRRK2-IN-1 is a type I inhibitor, which stabilizes LRRK2 in an active-like conformation^18^; while DARPin E11 enhances the stability of the LRRK2^RCKW^ fragment and facilitates cryo-EM analysis^56^. As expected, LRRK2^RCKW^ with LRRK2-IN-1 adopted an active-like conformation (**Fig. 3a-b**). LRRK2^RCKW^–DARPin E11 complexes were captured in three distinct oligomerization states: monomer, dimer and tetramer, of which the tetramer with *D*2 symmetry is likely representing a DARPin E11-induced artefact (**Extended Data Fig. 4a**). In contrast to the inactive full-length structure, Rab10 was only seen bound to site 4 of LRRK2^RCKW^ in the monomeric state and is incompatible with the dimeric active-like state of LRRK2^RCKW^ (**Extended Data Fig. 4a**).

**Fig. 3.**
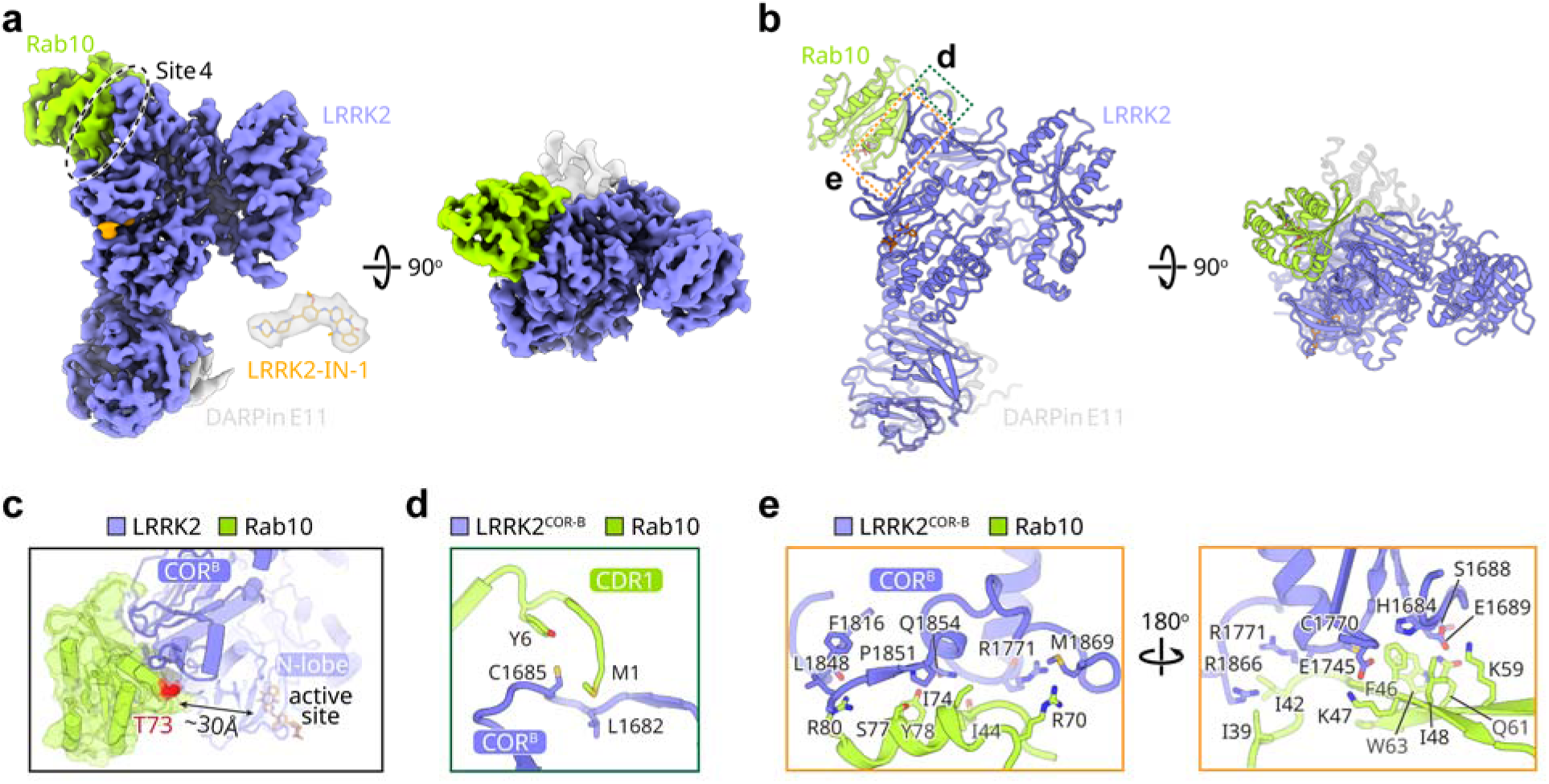
| Structure of active-like LRRK2^RCKW^ in complex with Rab10. **a-b**, Cryo-EM structure of active-like LRRK2^RCKW^ in complex with Rab10, shown in two views. Color code: Rab10, limon; LRRK2, slate; DARPin E11, grey. Density for the LRRK2-IN-1 inhibitor is shown. LRRK2 site 4 engaged by Rab10 is indicated with a dashed oval. **c,** Zoom-in view of the Rab10–LRRK2 interaction. Rab10-interacting domains in LRRK2 are labeled. The distance between the Rab10 Thr73 site and the LRRK2 active site is indicated. **d-e,** Interfaces between Rab10 (CDR1 region, **d**; Switch I–Interswitch–Switch II motifs, **e**) and the LRRK2 COR-B subdomain. Side chains of interface residues are shown as sticks.

The interactions between Rab10 with active-like LRRK2^RCKW^ are similar to those observed with the inactive monomer (**Fig. 1a-d, 3c**). Specifically, Rab10 CDR1 and Switch I-Interswitch-Switch II motifs interact with COR-B subdomain of LRRK2^RCKW^ (**Fig. 3c-e**). The N-terminus of Rab10 interacts with LRRK2^RCKW^, but in a different manner compared to the inactive monomer—in the active state, Rab10 Met1 and Tyr6 form hydrophobic interactions with COR-B residues Leu1682 and Cys1685, respectively (**Fig. 3d**); while in the inactive complex, Rab10 Met1 engages with Pro1661 and Ile1662 from COR-A, and Lys3 forms a salt bridge with Glu1714 on COR-B (**Fig. 1c**). Interestingly, despite the active-like conformation, Rab10 Thr73 phosphorylation site remains positioned ∼30 Å away from the kinase active site in the Rab10–LRRK2^RCKW^ (**Fig. 3c**),

### Rab8A binding at site 4

We then determined the cryo-EM structure of the Rab8A–LRRK2 complex (**Extended Data Fig. 5a-d and Table S1**), employing the same experimental strategy used for Rab10. Given the high sequence similarity in their N-terminal regions of Rab10 and Rab8A, we expected that Rab8A would engage LRRK2 at site 4 in the same manner as Rab10. Indeed, Rab8A occupies the identical binding site at the LRRK2 dimer interface as Rab10 (**Fig. 4**), but there were also notable differences between the two complexes. First, Rab8A binds selectively to site 4 and shows no detectable occupancy at site 3 (**Extended Data Fig. 5a**). Second, Rab10–LRRK2 exists in both monomeric (2:1 stoichiometry) and dimeric forms (4:2 stoichiometry) (**Extended Data Fig. 1a**), whereas Rab8A–LRRK2 is exclusively dimeric (**Fig. 4a**), forming 2:2 or 1:2 assemblies (**Extended Data Fig. 5a**). The 1:2 species are likely to reflect lower binding affinity, as a three-fold molar excess of Rab8A was used for grid preparation. Neither monomeric Rab8A–LRRK2 complexes or free LRRK2 monomers were observed in cryo-EM grids, indicating that Rab8A binding promotes LRRK2 dimerization, a conformation that would disrupt 14-3-3 binding^52^. Third, Rab8A induces a counterclockwise rotation of the COR-B subdomain and the kinase N-lobe, in contrast to the clockwise rotation observed in the Rab10-bound structure (**Extended Data Fig. 5e**). In line with this distinct rotational trajectory, Rab8A does not promote the rearrangement of the kinase domain towards an intermediate state seen with Rab10. Instead, the kinase domain in the Rab8A–LRRK2 complex remains in the inactive conformation seen in Rab12–LRRK2 and previously reported inactive LRRK2 structures (**Extended Data Fig. 5g-i**)^16^. Lastly, Rab8A binding also induces opening of the LRRK2 dimer, resulting in an ∼15° relative rotation of one protomer with respect to the other similar in magnitude to that observed in the Rab10-bound complex (**Extended Data Fig. 5e**).

**Fig. 4.**
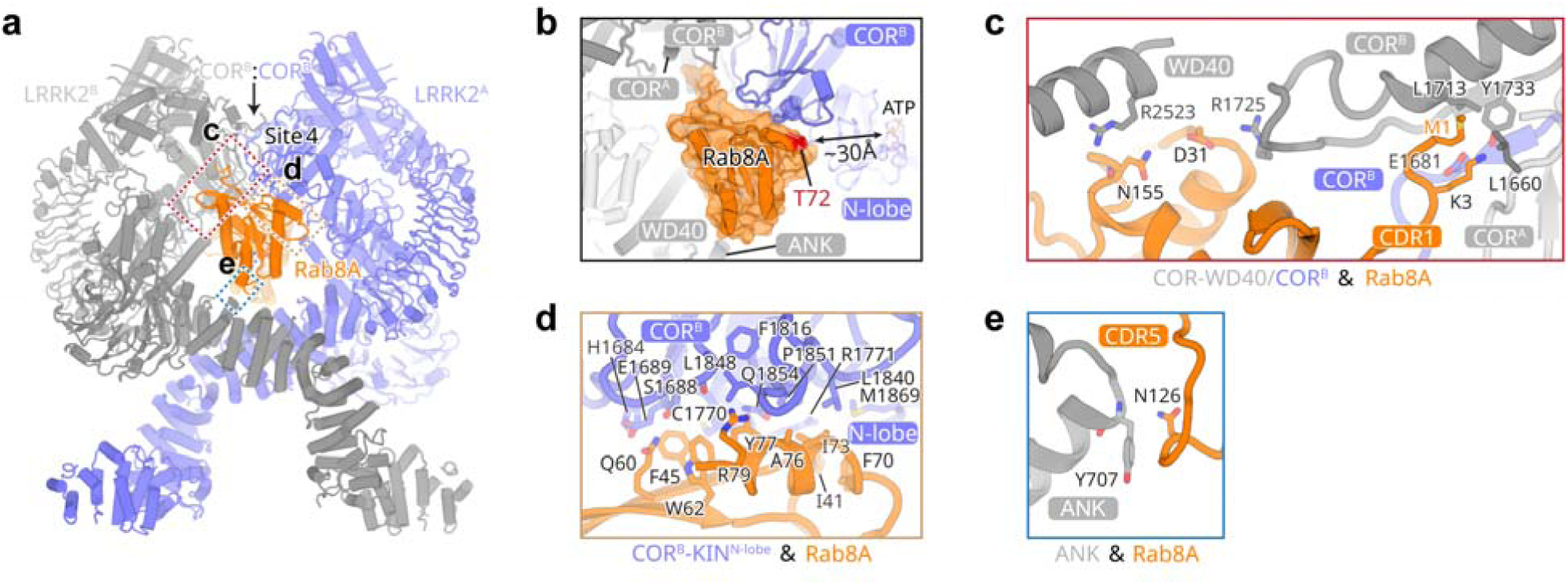
| Structural details of the Rab8A–LRRK2 complex. **a**, Cryo-EM structure of the Rab8A–LRRK2 complex in the LRRK2 dimer state. Color code: Rab8A, orange; LRRK2, slate and grey. LRRK2 site 4 engaged by Rab8A is indicated. **b,** Zoom-in view of the Rab8A–LRRK2 interaction. Rab8A-interacting domains in LRRK2 are labeled. The distance between the Rab8A Thr72 site and the LRRK2 active site is indicated. **c-e,** Interfaces between Rab8A and the LRRK2 COR/WD40 domains (**c**), the LRRK2 COR-B subdomain and KIN N-lobe (**d**), and the LRRK2 ANK domain (**e**). Side chains of interface residues are shown as sticks.

Our results revealed that binding of closely related Rab8A and Rab10 stabilizes distinct inactive conformations of the LRRK2 kinase domain. Rab8A binding keeps the LRKK2 kinase domain in a canonical inactive conformation, whereas Rab10 binding results in an intermediate configuration between inactive and active states (**Extended Data Fig. 5g, Fig. 2d**). Structural comparison of the Rab8A–LRRK2 and Rab10–LRRK2 complexes (**Extended Data Fig. 5j**) revealed the possible basis for this difference. Rab8A primarily engages the COR domain of LRRK2 protomer B via its CDR1 region. In contrast, Rab10’s CDR1 penetrates deeply into a hydrophobic pocket in the COR-B subdomain of LRRK2 protomer A, positioning Rab10 Thr5 and Tyr6 in close contact with its own Interswitch region. Additionally, Rab10’s Switch II region interacts closely with the kinase N-lobe, promoting a more closed kinase conformation. These distinct interaction modes might account for why Rab10, but not Rab8A, drives the kinase domain into the intermediate inactive state.

### The Rab-binding site 5 for Rab43 and site 6 for Rab5A

We next analyzed the interaction of LRRK2 and with two other Rab proteins, Rab43 and Rab5A, which belong to distinct Rab GTPase clades^57^. As mentioned, LRRK2 phosphorylates 14 Rab GTPases, including the Rab5 isoforms, although *in-vivo* Rab5 phosphorylation is not detected endogenously^25^, suggesting that it may be context-dependent or occur under specific physiological circumstances. In addition, LRRK2 binds two further Rab GTPases, Rab32 and Rab38. Phylogenetic analysis places these Rabs into four clades: i) Rab29/32/38, ii) Rab3A/B/C/D, Rab8A/B and Rab10/12/35, iii) Rab43, and iv) Rab5A/B/C (**Extended Data Fig. 3**)^57^. Rab43, a regulator of GPCR membrane trafficking^58, 59^, is sequence-divergent from the rest, whereas Rab5A (as well as Rab5B/C) regulates vesicular transport from the plasma membrane to early endosomes^60^.

We resolved the Rab43–LRRK2 complex at an overall resolution of 4.3 Å (**Extended Data Fig. 6a-d and Table S1**). The complex adopts a monomeric state with a 1:1 stoichiometry between Rab43 and LRRK2 (**Extended Data Fig. 6a**). Rab43 binds a previously unidentified binding site on LRRK2, which we designated site 5, located at the ARM–ANK domain interface (**Fig. 5a-b**). The interaction involves the ARM13–15 motifs and ANK domain from LRRK2 and Rab43 Switch I–Interswitch–Switch II regions, with key interface residues conserved among all LRRK2 Rab substrates (**Extended Data Fig. 3**). Compared to other structurally characterized Rab–LRRK2 complexes (Rab29, Rab12, Rab8A, Rab10); Rab43 has fewer contacts with LRRK2. This reduced interaction surface is consistent with the apparent weaker affinity of Rab43, as ∼50% of LRRK2 monomers remained unbound, despite using a five-fold molar excess of Rab43 during grid preparation (**Extended Data Fig. 6a**).

**Fig. 5.**
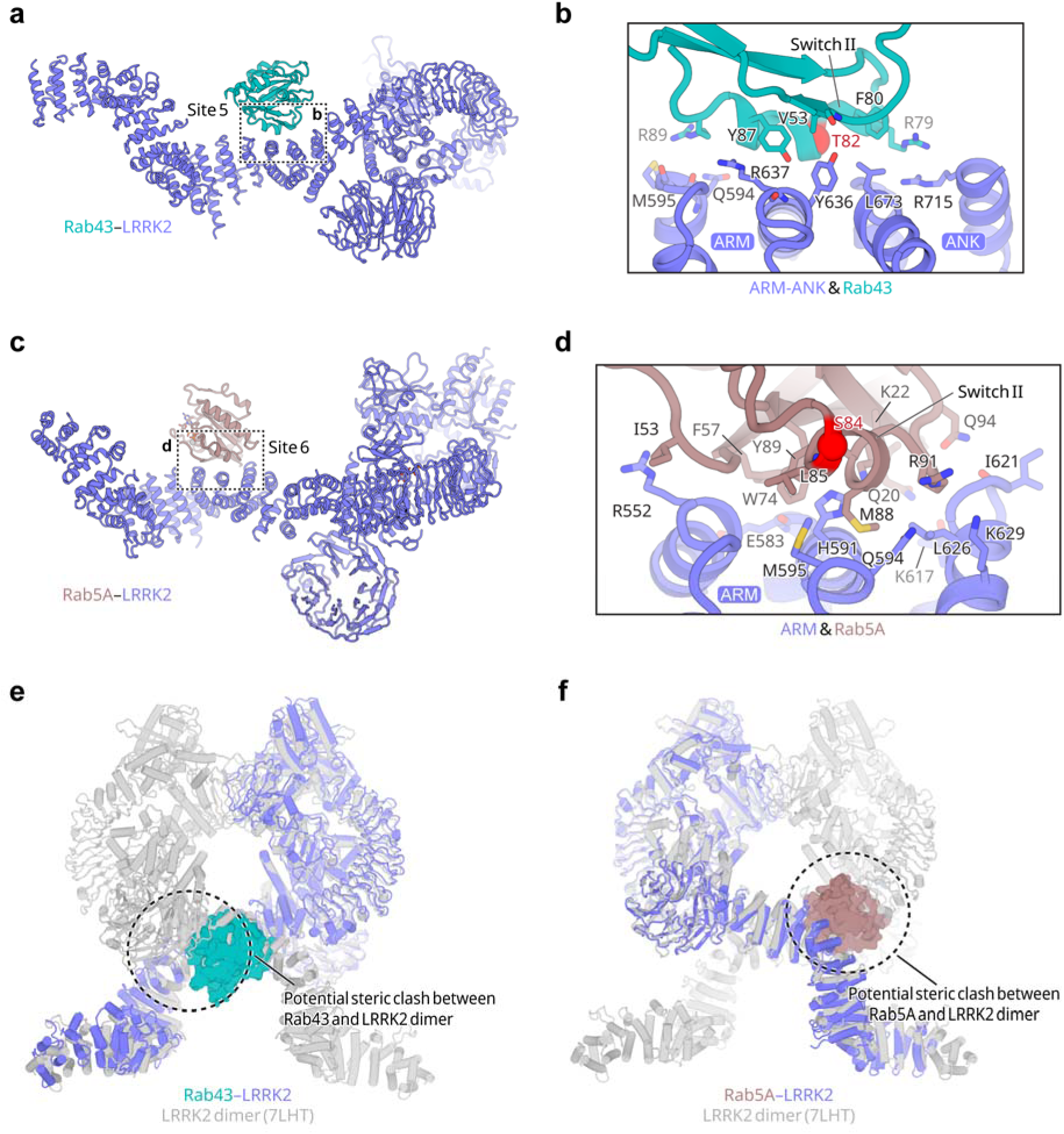
| Structural analysis of the Rab43–LRRK2 and Rab5A−LRRK2 complexes. **a**, Cryo-EM structure of the Rab43–LRRK2 complex. Color code: LRRK2, slate; Rab43, teal. **b,** Zoom-in view of the Rab43−LRRK2 interface. The side chains of interface residues are shown as sticks. The Rab43 phosphorylation site Thr82 within switch II motif is marked with a red sphere. **c,** Cryo-EM structure of the Rab5A−LRRK2 complex. Color code: LRRK2, slate; Rab5A, dirtyviolet. **d,** Zoom-in view of the Rab5A−LRRK2 interface. The side chains of interface residues are shown as sticks. The Rab5A phosphorylation site Ser84 within switch II motif is marked with a red sphere. **e,** Comparison of the Rab43–LRRK2 complex with the LRRK2-alone dimer (PDB 7LHT). The potential steric clash between Rab43 and LRRK2 dimer is indicated in a dashed circle. **f,** Alignment of the Rab5A−LRRK2 complex with the LRRK2-alone dimer (PDB 7LHT). The potential steric clash between Rab5A and LRRK2 dimer is indicated in a dashed circle.

We determined the cryo-EM structure of the Rab5A−LRRK2 complex at an overall resolution of 3.8 Å (**Extended Data Fig. 6e-h; Table S1**). Similarly, Rab5A−LRRK2 complexes adopt a monomeric 1:1 stoichiometry (**Extended Data Fig. 6e**). This structure revealed another distinct binding site, site 6, located within the ARM domain, adjacent to site 5 (**Fig. 5c-d**). The interface involves conserved Rab5A Switch I–Interswitch–Switch II residues (**Extended Data Fig. 3**), together with the LRRK2 ARM12–14 motifs.

In both complexes, the phosphorylation sites, Rab43-Thr82 and Rab5A-Ser84, are positioned far from the LRRK2 kinase active site, suggesting that the primary role of sites 5 and 6 function primarily in substrate recruitment/docking. Binding of either Rab43 or Rab5A introduces steric clashes with the opposing protomer in an LRRK2 dimer (**Fig. 5e-f**), accounting for the exclusive monomeric conformation observed for both complexes.

### A systematic analysis of LRRK2-Rab GTPases

By combining our new structures with prior work, we revealed six distinct Rab-binding sites on LRRK2. To assess whether these represent the complete repertoire of Rab-interaction sites, we integrated data from AlphaFold3 (AF3) modelling predictions^61^, sequence conservation analyses, and our cryo-EM structures. This enabled us to systematically evaluate all known LRRK2-interacting Rabs, including 14 substrates and 2 binders (Rab32/38) and assigned them to one of the six defined sites. (**Fig. 6, Extended Data Fig. 7**)^25, 26, 33^.

**Fig. 6.**
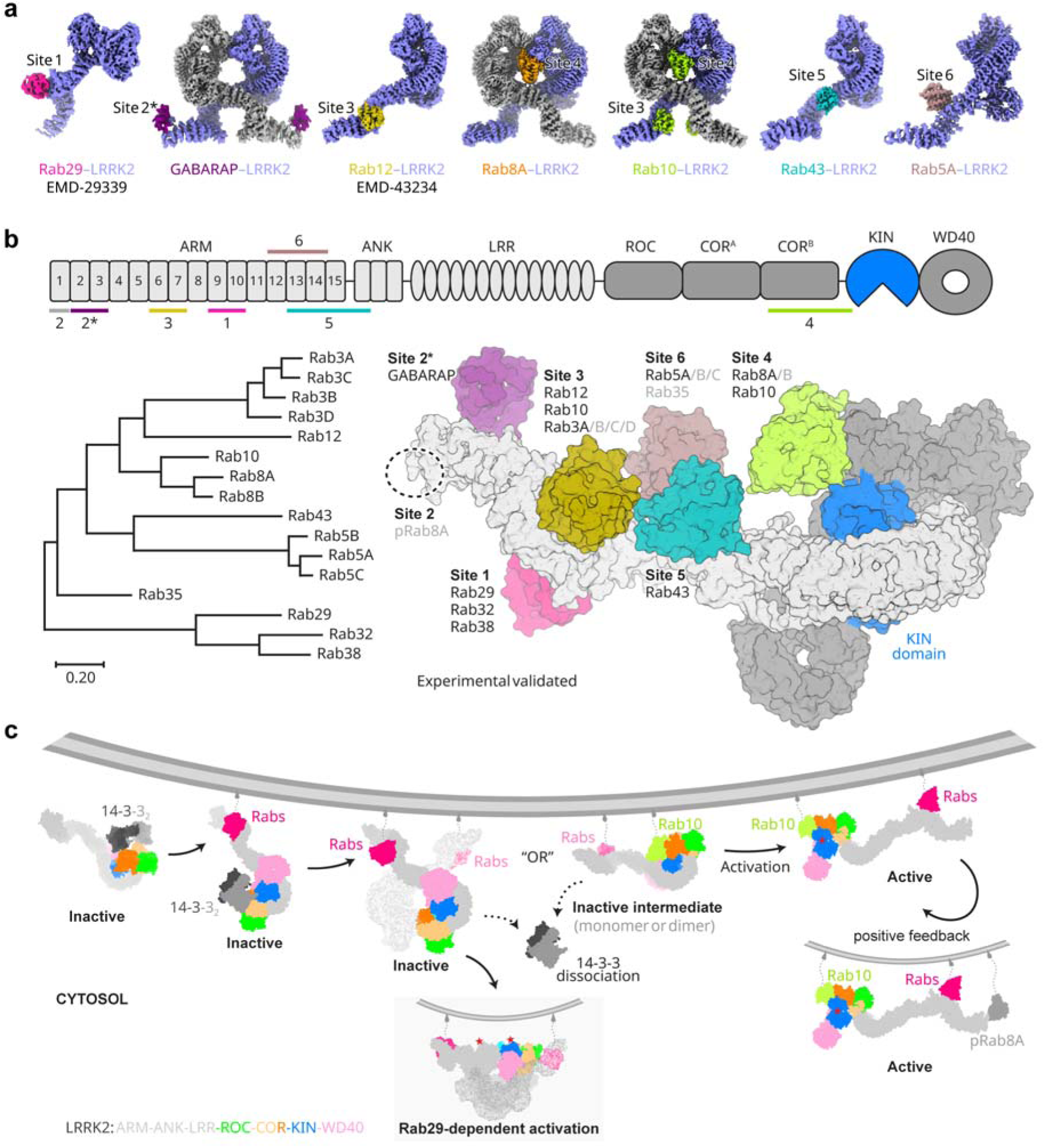
| Systematic structural analysis of LRRK2 bound to substrate or non-substrate Rab GTPases and a proposed model of LRRK2 activation. **a**, Cryo-EM structures of LRRK2 in complex with Rab29, GABARAP, Rab12, Rab8A, Rab10, Rab43, and Rab5A. Rab binding sites on LRRK2 are indicated. Color code: LRRK2 (slate/grey); Rab29 (magenta); GABARAP (purple); Rab12 (olive); Rab8A (orange); Rab10 (limon); Rab43 (teal); Rab5A (dirtyviolet). **b,** Distinct Rab-binding sites on LRRK2. Top: domain architecture of LRRK2 annotated with experimental identified Rab-binding sites. Bottom left: Maximum likelihood phylogenetic tree of substrate and non-substrate Rab GTPases of LRRK2. Bottom right: Structural model of LRRK2 displaying experimentally validated Rab binding sites and possible sites for those currently lacking structural data. Site 2* indicates the GABARAP-binding site in proximity to site 2 for pRab8A binding. The LRRK2 KIN domain is colored blue. **c,** Proposed model for LRRK2 membrane recruitment and activation. In the cytosol, LRRK2 is maintained in an inactive state by binding to the 14-3-3 scaffold protein. Activation is tightly linked to organelle membranes, where Rab GTPases recruit the LRRK2–14-3-3 complex. Upon membrane recruitment, concentration-dependent LRRK2 dimerization or docking of a second Rab at site 4 triggers 14-3-3 dissociation. This is followed by major conformational changes—driven by signals like asymmetric tetramerization *via* Rab29 or GTP binding at the ROC domain—that expose the kinase (KIN) domain for substrate phosphorylation. Finally, phosphorylated Rab8 could serve as positive feedback, further enhancing LRRK2 kinase activity.

We had previously established that site 1 binds Rab29 and Rab32^17^. Based on AF3 predictions and sequence similarity, Rab38 should occupy same site. To test this, we obtained a low-resolution cryo-EM structure of the Rab38–LRRK2 complex (**Extended Data Fig. 9a-c**) that confirmed Rab38 occupancy at site 1. Site 2 was associated with binding of phosphorylated Rab8A based on biochemistry and cellular evidence^42^, but the detailed structure remains elusive. AF3 models using either full-length LRRK2 or its ARM domain with pRab8A resulted in low-confidence predictions that didn’t align well with the biochemical data (**Extended Data Fig. 7**). Recent studies have identified a STING−CASM−GABARAP pathway, in which GABRAP recruites and activates LRRK2 at stressed lysosomes. The GABAPAP binding site was modelled and is close to site 2^51, 62^. We determined the cryo-EM structure of the dimeric GABARAP−LRRK2 complex at a resolution of 4.0 Å, which confirmed the binding mode of GABARAP (**Extended Data Fig. 10; Fig. 6a-b**)^51^. We assign Rab3A/B/C/D, Rab10, and Rab12 to site 3. Cryo-EM data of the Rab3A/10/12–LRRK2 complexes (**Fig. 1a; Extended Data Fig. 9d-f**) together with high sequence conservation among Rab3 subfamily support this assignment. Of note, AF3 prediction places Rab3A and its close homolog Rab3D bound at a site different from site 3, contradictory to our experimental observation (**Extended Data Figs. 7 and 9d-f**). Site 4, located at the LRRK2 dimer interface, engages Rab8A and Rab10 as predicted by AF3 when correct Rab:LRRK2 stoichiometry is specified. Given its high homology to Rab10 and Rab8A, Rab8B is predicted to interact in a similar manner. Site 5 is occupied by Rab43, whereas Site 6 accommodates Rab5A—and likely Rab5B/C—as evidenced by both cryo-EM and supported by AF3 modelling. AF3 also predicts that Rab35 engages with LRRK2 through site 6 (**Extended Data Figs. 7**).

AF3 predictions for most Rab–LRRK2 complexes, except Rab29 and pRab8A, yielded low ipTM scores (<0.6), likely explaining substantial discrepancies observed in some Rab–LRRK2 complexes (e.g., Rab3A and Rab32; **Extended Data Fig. 8a-b**). Even when binding sites on LRRK2 were correctly identified, the predicted interactions showed substantial differences with cryo-EM data, as exemplified in the case of Rab43 and Rab5A (**Extended Data Fig. 8c-d**). There were also deviations from the experimental data regarding LRRK2 conformation or interface geometry (e.g., Rab10 and Rab8A; **Extended Data Fig. 8e-f**), even when the correct stoichiometry was specified in the input. These observations highlight the necessity of experimental validation for accurately delineating Rab–LRRK2 interactions that are important in guiding subsequent biochemical characterization^63^.

## Discussion

Here we elucidated the molecular interactions between LRRK2 and Rab GTPases, establishing six distinct Rab-binding sites on LRRK2 (**Fig. 6a-b**), including three previously unidentified ones. Site 4, occupied by Rab10 and Rab8A, is of particular interest. Rab10 and Rab8A are two most abundant Rab GTPases in mouse embryonic fibroblasts (MEF), with reported ∼1.25 million and ∼0.65 million copies per cell, respectively, compared to ∼6,000 copies of LRRK2 per cell. Binding of Rab8A or Rab10 at site 4 promotes LRRK2 dimerization and may facilitate dissociation of 14-3-3 proteins, thereby contributing to kinase activation. In addition, Rab10 engagement stabilized the kinase domain in an intermediate conformation with structural features indicating it is progressing towards to an active state. Accordingly, some mutations disrupting site 4 reduced LRRK2 kinase activity. Finally, with respect to membrane geometry, site 4 is positioned similarly to sites 1 and 3, both of which are involved in membrane recruitment of LRRK2. Collectively, these observations support a model in which LRRK2 activation may involve multi-step recruitment, with site 4 (or other sites) serving as a secondary docking site in addition to Rab12 or Rab29/32/38.

We illustrated one such scenario in **Fig. 6c**. Upon stimulation, cytosolic LRRK2, likely in complex with 14-3-3, is recruited to cellular membrane by Rab GTPases. This recruitment may occur through multiple pathways, including lysosomal damage-induced and Rab12-dependent pathways *via* site 3; Parkinson’s disease mutation-associated processes mediated by Rab29 or Rab32 through site 1; or potentially as-yet-unidentified mechanisms involving other sites. This membrane recruitment is followed by dissociation of 14-3-3 and subsequent dephosphorylation of LRRK2 pS935. These events could be triggered by increased local concentration at membranes and subsequent LRRK2 dimerization of LRRK2 or by secondary docking of Rab at site 4. LRRK2 then undergoes conformational rearrangements that exposes the kinase domain for substrate phosphorylation. Lastly, phosphorylated Rab8 could bind to site 2, providing a positive feedback pathway to further activate LRRK2^42^. In the context of Rab29, LRRK2 activation could also be achieved through asymmetric tetramerization in a Rab29– and LRRK2-concentration dependent manner^17^.

The identification and structural characterization of Rab-binding sites 4-6 open new avenues to dissect LRRK2 roles in diverse biological contexts. For example, the physiological functions of Rab5A and Rab43 in LRRK2-mediated signaling remain poorly defined. Our structural data provided a precise molecular framework to design mutations that specifically disrupt these interactions, without broadly perturbing LRRK2 architecture. Such targeted approaches will enable mechanistic interrogation of how Rab5A and Rab43-dependent pathways contribute to LRRK2 biology and disease-relevant processes.

It is noteworthy that, across all Rab–LRRK2 complexes determined to date, whether inactive or active-like conformations, the Rab substrates were positioned too distantly from the kinase active site to permit direct phosphate transfer from ATP. This spatial separation raises a key question regarding how LRRK2 achieves selective substrate recognition and phosphorylation. We propose two non-mutually exclusive possibilities. First, substrate specificity could be governed by the spatial proximity conferred through Rab docking at sites 1–6. In this model, initial engagement at these docking sites concentrates and orients specific Rab GTPases near LRRK2, with subsequent conformational rearrangements enabling productive phosphorylation. Thus, selective recognition would arise from the combined effects of docking geometry, membrane localization, and local concentration. Second, an additional as-yet-unidentified Rab-binding site adjacent to the kinase active site may exist. This site could be of low-affinity and be transiently occupied, rendering it difficult to capture for biochemical and cryo-EM analysis. Transient engagement at this catalytic-proximal site might occur only after prior docking at one of the defined sites and conformational activation of the kinase domain.

In closing, the present study established a foundational framework for biochemical characterization of Rab–LRRK2 signaling and for the development of pharmacological tools and therapeutics aimed at modulating these Rab–LRRK2 interactions.

## Methods

### Expression and purification of human full-length LRRK2 and LRRK2^RCKW^

Full-length human LRRK2 and the LRRK2^RCKW^ construct were expressed and purified as previously described^16, 18^. An N-terminal GFP tag followed by a PreScission protease cleavage site was engineered into each construct and cloned into a BacMam expression vector. Bacmids were generated in *E. coli* DH10Bac cells (Invitrogen), and recombinant baculoviruses were produced by transfecting Sf9 cells using Cellfectin^TM^ II (Gibco). Following two rounds of amplification in Sf9 cells, the viruses were used to transduce HEK293S GnTI^─^ cells for protein expression. When the HEK293S GnTI^─^ cells reached a density of ∼2−3×10^6^ cells per mL in Expi293^TM^ expression medium (Gibco) supplemented with 2% FBS (Gibco), baculoviruses were added at 10% (v/v). Infected cells were incubated overnight at 37 °C, after which protein expression was induced with 10 mM sodium butyrate. Cells were then cultured at 30 °C for an additional 48−60 h prior to harvest.

For full-length LRRK2, cell pellet from 600 mL culture was resuspended in 30 ml lysis buffer (20 mM Tris-HCl pH 8.0, 200 mM NaCl, 5% glycerol, 2 mM DTT, 1 mM PMSF, and Pierce Protease Inhibitor Tablets) and lysed via brief sonication. The lysate was cleared by high-speed centrifugation (39,800 × *g* for 1 h), and the supernatant was incubated with 1 mL of CNBr-activated Sepharose beads (Cytiva) coupled to 1 mg of high-affinity GFP nanobodies (GFP-NB)^64^. The GFP tag was removed by PreScission protease cleavage at 4 °C. LRRK2 was further purified by size-exclusion chromatography (SEC) using a Superose 6 Increase 10/300 GL column (Cytiva) equilibrated in 20 mM Tris-HCl pH 8.0, 200 mM NaCl, and 2 mM DTT. The peak fractions were collected and concentrated to 10 mg/ml (as determined by OD_280_) using a 100-kDa MWCO centrifugal concentrator (Amicon), flash-frozen in liquid nitrogen, and stored at −80 °C.

For LRRK2^RCKW^, cell pellet was resuspended in lysis buffer (20 mM HEPES pH 7.4, 300 mM NaCl, 5% glycerol, 2 mM DTT, 1 mM PMSF, and Pierce Protease Inhibitor Tablets) and lysed by sonication. After centrifugation at 39,800 × *g* for 1 h, the protein was captured using 1 mL of GFP-NB−coupled Sepharose beads. To remove heat shock protein contaminants, the beads were washed with lysis buffer supplemented with 1 mM MgCl_2_ and 2 mM ATP. Following PreScission protease cleavage at 4 °C, the protein was eluted and concentrated to 2 mg/mL (OD_280_), flash-frozen in liquid nitrogen, and stored at −80 °C.

### Purification and assembly of the LRRK2^RCKW^−DARPin E11 complex

The synthesized cDNA encoding N-terminal His_8_-tagged DARPin E11^56^ was subcloned into pET-15b plasmid (Sigma). Recombinant proteins were overexpressed in *E. coli* strain BL21 (DE3) using LB media supplemented with 0.05 mg/mL ampicillin. Cells were grown at 37 °C until OD_600_ reached 0.8. Protein expression was then induced with 0.4 mM IPTG, and the cells were cultured at 16 °C for an additional 20 h.

Cells were harvested and lysed via sonication in lysis buffer (20 mM Tris-HCl pH 8.0, 200 mM NaCl, 5% glycerol, and 1 mM PMSF). The cleared lysate was incubated with TALON Metal Affinity Resin (Takara), followed by washing with lysis buffer containing 10 mM imidazole. The protein was eluted with lysis buffer supplemented with 150 mM imidazole. Further purification was performed by SEC using a Superdex 200 Increase 10/300 GL column (Cytiva) equilibrated in storage buffer (20 mM Tris-HCl pH 8.0, 100 mM NaCl, and 2 mM DTT). Peak fractions were pooled, concentrated to 23 mg/mL (OD_280_), flash-frozen in liquid nitrogen, and stored at −80 °C.

To assemble the LRRK2^RCKW^–DARPin E11 complex, purified LRRK2^RCKW^ and DARPin E11 were mixed at a 1:3 molar ratio. The complex was isolated via SEC on a Superose 6 Increase 10/300 GL column equilibrated in 20 mM HEPES pH 7.4, 200 mM NaCl, and 2 mM DTT. The final purified complex was concentrated to 8 mg/mL, flash-frozen, and stored at −80°C.

### Expression and purification of Rab GTPases and GABARAP

DNA sequences encoding the human GTP-locked mutants Rab8A (Q67L, residues 1−176), Rab10 (Q68L, residues 1−181), Rab43 (Q77L, residues 1−190), Rab5A (Q79L, residues 15−184), Rab3A (Q81L, residues 1−199), Rab38 (Q69L, residues 1−181), and wild-type GABARAP were synthesized. The cDNA of Rab8A, Rab10, Rab43, Rab3A, and GABARAP was subcloned into the pET-15b vector (Sigma), which attaches a His_6_ tag and a thrombin cleavage site at the N-terminus. Rab5A and Rab38 cDNA were subcloned into a modified pGEX vector (Sigma), which attaches a GST tag and a TEV protease cleavage site at the N-terminus. All recombinant proteins were overexpressed in *E. coli* strain BL21 (DE3) cells grown in LB media supplemented with 0.05 mg/mL ampicillin at 37 °C until the OD_600_ reached 0.8. Protein expression was induced with 0.4 mM IPTG, followed by incubation at 16 °C for 20 h. Cells were then harvested, flash-frozen in liquid nitrogen, and stored at −80 °C.

Rab5A and Rab38 were purified as previously described^19^. Cells were harvested and lysed via sonication in lysis buffer (20 mM Tris-HCl pH 8.0, 200 mM NaCl, 5% glycerol, 5 mM MgCl_2_, and 1 mM PMSF). The resulting lysates were cleared by centrifugation at 38,000 × *g* for 45 min. Proteins were captured using GST affinity chromatography (Glutathione Sepharose beads, Cytiva) in lysis buffer and eluted in lysis buffer supplemented with 20 mM GSH. To ensure nucleotide loading, the eluates were incubated with 1 mM GTP at 4 °C overnight. Final purification was performed via SEC using a Superdex 200 Increase 10/300 GL column (Cytiva) in storage buffer (20 mM Tris-HCl pH 8.0, 100 mM NaCl, 1 mM MgCl_2_, and 2 mM DTT). For structural studies, the GST tag was removed by on-column cleavage with TEV protease overnight at 4 °C prior to SEC. Peak fractions were pooled, concentrated to approximately 15 mg/mL (OD_280_), flash-frozen in liquid nitrogen, and stored at −80 °C.

Cells expressing Rab8A, Rab10, Rab43, and Rab3A were resuspended in lysis buffer (20 mM Tris-HCl pH 8.0, 200 mM NaCl, 5% glycerol, 5 mM MgCl_2_, and 1 mM PMSF) and lysed via sonication. Cleared lysates were incubated with TALON Metal Affinity Resin (Takara) and washed with lysis buffer supplemented with 10 mM imidazole. To ensure nucleotide loading, the resin was incubated with 1 mM GppNHp overnight at 4 °C. The protein was subsequently eluted using lysis buffer containing 150 mM imidazole and further purified by SEC (Superdex 200 Increase 10/300 GL, Cytiva) in storage buffer (20 mM Tris-HCl pH 8.0, 100 mM NaCl, 1 mM MgCl_2_, and 2 mM DTT). Peak fractions were pooled, concentrated to approximately 15 mg/mL (OD_280_), flash-frozen in liquid nitrogen, and stored at −80 °C.

GABARAP cell pellets were resuspended and lysed by sonication in lysis buffer (20 mM Tris-HCl pH 8.0, 200 mM NaCl, 5% glycerol, and 1 mM PMSF). The cleared lysates were incubated with TALON Metal Affinity Resin (Takara), followed by washing with lysis buffer supplemented with 10 mM imidazole. The protein was eluted using lysis buffer containing 150 mM imidazole and further purified via SEC on a Superdex 200 Increase 10/300 GL column (Cytiva) equilibrated in storage buffer (20 mM Tris-HCl pH 8.0, 100 mM NaCl, 1 mM MgCl_2_, and 2 mM DTT). Peak fractions were pooled and concentrated to 13.6 mg/mL (OD_280_), flash-frozen in liquid nitrogen, and stored at −80 °C.

### Cryo-EM sample preparation for full-length LRRK2 in complex with Rab proteins

Purified full-length LRRK2 was incubated with Rab3A, Rab5A, Rab8A, Rab10, Rab38, or Rab43 on ice for 1 h. Final concentration were 30 μM for LRRK2 and 90 μM to 150 μM for the Rab proteins in a buffer containing 2 mM ATP and 1 mM MgCl_2_. 2.3 mM fluorinated Fos-Choline-8 was added right before freezing the grids, and then 3.5 μL of protein sample was applied to a glow-discharged Quantifoil R1.2/1.3 300 Au holey carbon grid (Quantifoil). Grid was incubated in a FEI Vitrobot IV chamber (16 °C, 95% relative humidity) for 20 s, blotted for 3.5 s with filter paper 595 (blot force −3), and vitrified by plunging into liquid nitrogen-cooled ethane.

### Cryo-EM sample preparation for Rab10–LRRK2^RCKW^–DARPin E11 complex

Purified LRRK2^RCKW^–DARPin E11 complex and Rab10 were incubated on ice for 1 h at final concentration of 40 μM and 90 μM, respectively (1:2.3 molar ratio, in the presence of 1 mM MgCl_2_. Subsequently, 100 μM LRRK2-IN-1 was added, followed by incubation on ice for 15 min and centrifugation at 4 °C for 10 min. 2.3 mM fluorinated Fos-Choline-8 was added right before vitrification, and then 3.5 μL of protein sample was applied to a glow-discharged Quantifoil R1.2/1.3 300 Au holey carbon grid (Quantifoil). Grid was incubated in a FEI Vitrobot IV chamber (16 °C, 95% relative humidity) for 20 s, blotted for 3.5 s with filter paper 595 (blot force −3), and vitrified by plunging into liquid nitrogen-cooled ethane.

### Cryo-EM sample preparation for GABARAP–LRRK2 complex

Purified full-length LRRK2 and GABARAP were incubated on ice for 1 h at final concentrations of 35 μM and 115 μM, respectively (1:3.3 molar ratio), in the presence of 2 mM ATP and 1 mM MgCl_2_. 2.3 mM fluorinated Fos-Choline-8 was added to the mixture prior to grid preparing, and then 3.5 μL of protein sample was applied to a glow-discharged Quantifoil R1.2/1.3 300 Au holey carbon grid (Quantifoil). Grid was incubated in a FEI Vitrobot IV chamber (16 °C, 95% relative humidity) for 20 s, blotted for 5.0 s with filter paper 595 (blot force 0), and vitrified by plunging into liquid nitrogen-cooled ethane.

### Cryo-EM data acquisition

The Rab8A–LRRK2 and Rab10−LRRK2 datasets were collected on a Titan Krios (Thermo Fisher Scientific) transmission electron microscope equipped with a K3 direct electron detector and post-column GIF energy filter (Gatan). Data collection was performed in an automated manner using EPU (Thermo Fisher Scientific). Movies of Rab8A−LRRK2 were recorded at defocus values from −1.0 to −2.6 μm at a magnification of 81,000 in hardware binning mode, corresponding to a pixel size of 1.057 Å at the specimen. During 4.0 s exposure, 50 frames were collected with a total electron dose of 50.2 e-/Å^2^ (at a dose rate of 1.004 e-/frame/Å^2^). Movies of Rab10−LRRK2 were recorded at defocus values from −0.8 to −2.4 μm at a magnification of 130,000 in hardware binning mode, corresponding to a pixel size of 0.6485 Å at the specimen. During 1.8 s exposure, 60 frames were collected with a total electron dose of 60.2 e-/Å^2^ (at a dose rate of 1.0 e-/frame/Å^2^).

The Rab43–LRRK2 and Rab3A−LRRK2 datasets were collected on a Titan Krios (Thermo Fisher Scientific) transmission electron microscope equipped with a K3 direct electron detector and post-column GIF energy filter (Gatan). Data collection was performed using SerialEM software^65^. Movies of Rab43−LRRK2 were recorded at defocus values from −1.0 to −2.8 μm at a magnification of 81,000 in hardware binning mode, corresponding to a pixel size of 1.105 Å at the specimen. During 5.2 s exposure, 50 frames were collected with a total electron dose of 50.8 e-/Å^2^ (at a dose rate of 1.02 e-/frame/Å^2^). Movies of Rab3A−LRRK2 were recorded at defocus values from −1.0 to −2.8 μm at a magnification of 81,000 in hardware binning mode, corresponding to a pixel size of 1.105 Å at the specimen. During 3.8 s exposure, 60 frames were collected with a total electron dose of 61.0 e-/Å^2^ (at a dose rate of 1.02 e-/frame/Å^2^).

The Rab10–LRRK2^RCKW^–DARPin E11, Rab5A–LRRK2, GABARAP–LRRK2, and Rab38−LRRK2 datasets were collected on a Talos Arctica (Thermo Fisher Scientific) operated at 200 kV, equipped with a K3 Summit direct electron detector and a post-column GIF energy filter (Gatan). Automated data collection was performed using EPU (Thermo Fisher Scientific). Movies of Rab10–LRRK2^RCKW^–DARPin E11 were recorded at defocus values from −1.2 to −3.0 μm at a magnification of 130,000, which corresponded to a pixel size of 0.63 Å at the specimen level. During 1.7 s exposure, 60 frames were collected, corresponding to a total electron dose of 61.2 e-/Å^2^ (at a dose rate of 1.02 e-/frame/Å^2^). Movies of Rab5A–LRRK2 and GABARAP–LRRK2 were recorded at defocus values from −1.2 to −2.8 μm at a magnification of 130,000, which corresponded to a pixel size of 0.63 Å at the specimen level. During 2.0 s exposure, 60 frames were collected, corresponding to a total electron dose of 60 e-/Å^2^ (at a dose rate of 1.0 e-/frame/Å^2^). Movies of Rab38–LRRK2 were recorded at defocus values from −0.8 to −2.4 μm at a magnification of 79,000, which corresponded to a pixel size of 1.044 Å at the specimen level. During 4.0 s exposure, 60 frames were collected, corresponding to a total electron dose of 61.3 e-/Å^2^ (at a dose rate of 1.02 e-/frame/Å^2^).

### Data processing for Rab10−LRRK2 complex

For the Rab10−LRRK2 complex, 22,961 movies were collected and aligned using the MotionCor2 dose-weighted alignment option^66^. Contrast transfer function (CTF) estimation was performed via CTFFIND4^67^. Micrographs with a CTF fit worse than 6.0−6.5 Å (as determined by CTFFIND4) were discarded from further processing (with 21,130 good images remaining). Particles were selected using the cryoSPARC Blob picker^68^, followed by multiple rounds of 2D classification to remove ice artifacts, carbon edges, and false-positive particles containing noise. Two distinct classes, corresponding to monomeric and dimeric Rab10−LRRK2, were identified during 2D classification. Following *ab initio* reconstruction and heterogeneous refinement, 43,982 particles were assigned to the monomeric class and 113,857 particles to the dimeric class. For the monomer, non-uniform (NU) refinement yielded a 4.1 Å map. For the dimer, NU refinement with *C*2 symmetry and CTF refinement produced a 3.3 Å map. Symmetry expansion and focused 3D refinement were subsequently utilized to improve the map quality of the LRRK2 N-terminal ARM domain with Rab10 binding. All resolution estimates were calculated using the gold-standard Fourier shell correlation (FSC = 0.143)^69^ criterion, and local resolution was estimated within cryoSPARC. The density maps were B-factor sharpened for model building and figure generation.

### Data processing for Rab8A−LRRK2 complex

For the Rab8A–LRRK2 complex, 18,601 movies were collected and preprocessed as previously described, with 18,109 good images retained. Particles were selected using the cryoSPARC Blob picker^68^, followed by several rounds of 2D classification. Two distinct classes were observed, corresponding to Rab8A–LRRK2 dimers with stoichiometries of 2:2 and 1:2 (Rab8A:LRRK2), respectively. Following *ab initio* reconstruction, heterogeneous refinement was performed to partition the dataset, resulting in 564,974 particles for the 2:2 state and 552,037 particles for the 1:2 state. For the 1:2 stoichiometry, NU refinement with *C*1 symmetry yielded a 3.5 Å map. For the 2:2 stoichiometry, NU refinement with *C*2 symmetry and CTF refinement produced a 3.1 Å map. To further improve the density of the LRRK2 N-terminal ARM domain, we performed symmetry expansion followed by focused 3D refinement. All resolution estimates were determined using the gold-standard Fourier shell correlation (FSC = 0.143) criterion^69^, and local resolution was estimated within cryoSPARC. The density maps were B-factor sharpened for model building and figure generation.

### Data processing for Rab10−LRRK2^RCKW^−DARPin E11 complex

5,714 movies were collected for Rab10−LRRK2^RCKW^−DARPin E11 and preprocessed as previously described, with 5,245 good images retained. Particles were selected using the cryoSPARC Blob picker^68^, followed by several rounds of 2D classification. Three groups of good class were observed, corresponding to the monomeric, dimeric, and tetrameric states of LRRK2^RCKW^. Following *ab initio* reconstruction and heterogeneous refinement, 113,825 particles were assigned to the monomer, 31,418 to the dimer, and 32,248 to the tetramer. For the monomeric state, non-uniform (NU) refinement yielded a 3.7 Å map. For the dimeric and tetrameric states, NU refinement was performed by applying *C*2 and *D*2 symmetry, respectively, resulting in maps at 4.4 Å and 3.6 Å resolution. To further enhance map quality, symmetry expansion and focused 3D refinement were subsequently utilized. All resolution estimates were calculated according to the gold-standard Fourier shell correlation (FSC) using the 0.143 criterion^69^, and local resolution was estimated within cryoSPARC.

### Data processing for Rab43−LRRK2 complex

For the Rab43−LRRK2 complex, 8,112 movies were collected for Rab43−LRRK2 and preprocessed as previously described, with 7,971 good images remained. Particles were selected using the Blob picker in cryoSPARC^68^. Several rounds of the 2D classification were performed and two groups of good classes were observed, corresponding to LRRK2 monomeric and dimeric states. Following *ab initio* reconstruction and heterogeneous refinement, 131,849 particles were assigned to the monomeric class and 139,739 particles to the dimeric class (LRRK2 alone). To isolate the Rab43-bound population, the monomeric particles underwent further 3D classification, which yielded 59,498 particles for the Rab43–LRRK2 monomer. Subsequent NU refinement and CTF refinement produced a final map at 4.3 Å resolution. To improve the density of the C-terminal domains and the LRRK2 N-terminal ARM domain at the Rab43 binding interface, focused 3D refinement was performed. All resolution estimates were determined using the gold-standard Fourier shell correlation (FSC = 0.143) criterion^69^, and local resolution was estimated within cryoSPARC.

### Data processing for Rab5A−LRRK2 complex

For the Rab5A−LRRK2 complex, 5,623 movies were collected and preprocessed as previously described, with 5,386 good images remained. Particles were selected using the cryoSPARC Blob picker^68^, and following several rounds of 2D classification, a single population of good class was identified, corresponding to the Rab5A–LRRK2 monomer. *Ab initio* reconstruction and heterogeneous refinement resulted in 87,413 particles. Subsequent NU refinement and local CTF refinement yielded a final map at 3.8 Å resolution. To improve the density of the C-terminal domains and the LRRK2 N-terminal ARM domain at the Rab5A binding interface, focused 3D refinement was performed. The resolution estimate was calculated using the gold-standard Fourier shell correlation (FSC = 0.143) criterion^69^, and local resolution was estimated within cryoSPARC.

### Data processing for GABARAP−LRRK2 complex

For the GABARAP–LRRK2 complex, 10,174 movies were collected and preprocessed as previously described, with 9,689 good images retained. Particles were selected using the cryoSPARC Blob picker^68^, and several rounds of 2D classification were performed. A single population of good class was identified, corresponding to the GABARAP–LRRK2 dimer. *Ab initio* reconstruction and heterogeneous refinement resulted in 50,065 particles for the dimeric class. We then performed NU refinement with *C*2 symmetry and CTF refinement, yielding a final map at 4.0 Å resolution. To improve the density of the C-terminal domains and the LRRK2 N-terminal ARM domain at the GABARAP binding interface, symmetry expansion and focused 3D refinement were subsequently utilized. The resolution estimate was determined using the gold-standard Fourier shell correlation (FSC = 0.143) criterion^69^, and local resolution was estimated within cryoSPARC.

### Data processing for Rab38−LRRK2 and Rab3A−LRRK2 complexes

For the Rab38–LRRK2 and Rab3A–LRRK2 complexes, 2,256 and 2,583 movies were collected, respectively, and preprocessed as previously described. Particles were selected using the cryoSPARC Blob picker^68^, and several rounds of 2D classification revealed two populations of good classes for both complexes, corresponding to monomeric and dimeric states. Following *ab initio* reconstruction and heterogeneous refinement, 20,950 monomeric and 12,028 dimeric particles were identified for Rab38–LRRK2, while 12,746 monomeric and 17,591 dimeric particles were identified for Rab3A–LRRK2. NU refinement was performed for the monomers (*C*1 symmetry) and the dimers (*C*2 symmetry). For Rab38–LRRK2, the final maps reached resolutions of 6.3 Å (monomer) and 6.6 Å (dimer). For Rab3A–LRRK2, both the monomer and dimer reached a resolution of 7.3 Å. All resolution estimates were determined using the gold-standard Fourier shell correlation (FSC = 0.143) criterion^69^, and local resolution was estimated within cryoSPARC.

### Model building and refinement

For the Rab–LRRK2 complexes (Rab8A, Rab10, Rab43, Rab5A, and GABARAP), the reported structure of LRRK2 from the Rab12–LRRK2 complex (PDB: 8VH4) and AlphaFold3-predicted models of the respective Rabs were docked and manually adjusted into the cryo-EM maps using UCSF Chimera^70^ and Coot^71^. For the Rab10–LRRK2^RCKW^–DARPin E11 complex, the structural models of LRRK2^RCKW^ (PDB: 8FO7), DARPin E11 (PDB: 8U1B), and AlphaFold3-predicted Rab10 were similarly fitted. All structural models were refined against the density maps using the Phenix real-space refinement module^72^, with secondary structure and non-crystallographic symmetry (NCS) restraints applied. Model quality was assessed via Fourier shell correlation (FSC) curves calculated between the refined coordinates and the full maps, and geometries were validated using MolProbity^73^. All the figures were prepared in PyMOL (Schrödinger, LLC.), UCSF Chimera and UCSF ChimeraX^74^.

### Cell Culture, Transfection and Lysis

HEK293 (LRRK2 KO or Rab10 KO) cells were grown at 37 °C in Dulbecco’s modified Eagle medium supplemented with fetal calf serum (10%), *L*-glutamine (2 mM), penicillin (100 U/mL), and streptomycin (100 μg/mL). Cells were transfected in accordance to previously established protocols (https://dx.doi.org/10.17504/protocols.io.bw4bpgsn). Briefly, cells were cultured in 6 cm plates until 60−70% confluency. LRRK2 and Rab plasmids (1.6 μg LRRK2: 0.4 μg Rab) were incubated with PEI (3:1 PEI:DNA, pH 7.4, 15 min) in OPTI-MEM (100 μL per well) and introduced to the culture. After 24 h, the media was aspirated and lysis buffer (50 mM Tris-HCl pH 7.4, 10 mM Sodium β-glycerophosphate, 10 mM Sodium pyrophosphate, 150 mM NaCl, 10% w/v glycerol, complete EDTA-free protease inhibitor cocktail, PhosStop, 1% v/v Triton) was added. The lysates were centrifuged (17,000 x *g*, 10 min) and protein content in the supernatant was determined by Bradford assay. Lysates were stored at −20 °C.

### Immunoblotting

Immunoblotting of LRRK2 pathway components was undertaken using previously described methods (dx.doi.org/10.17504/protocols.io.ewov14znkvr2/v2). Briefly lysate samples were supplemented with LDS (1x) and BME (1.25% v/v), loaded into NuPage 4-12% Bis-Tris gels (Invitrogen), and run in MOPS buffer. Upon completion, the protein was transferred to an Amersham Protran 0.45 μm nitrocellulose membrane (Cytiva) by wet transfer in Tris-HCl (48 mM), glycine (39 mM), and MeOH (20% v/v) buffer. The membrane was cut, washed with TBST (20 mM Tris-HCl pH 7.5, 150 mM NaCl, 0.1% v/v Tween-20) and blocked with milk (5% w/v in TBST) for 1 h. After washing again, the membrane was incubated overnight with primary antibodies in BSA (5% w/v in TBST) at 4 °C. The treated membranes were washed and incubated with secondary antibodies (1:20,000) in milk (5% w/v in TBST) for 1 h, after which they were washed and imaged at 700 and 800 nm.

## Acknowledgments

We thank the support of the Cryo-Electron Microscopy Center at St. Jude Children’s Research Hospital and National University of Singapore, for their help with cryo-EM data collection; members of the Sun lab for helpful discussion; Dhayanitha Ranganathan Dhakshinamoorthy and Patricia Hixson for cell culture support. We thank the excellent technical support of the MRC Protein Phosphorylation and Ubiquitylation Unit (PPU) DNA cloning team (coordinated by Rachel Toth), sequencing service (coordinated by Gary Hunter), and tissue culture team (coordinated by Edwin Allen).

## Author Contributions

HZ and JS designed the project. HZ performed sample preparation and structural analysis. LE designed, performed and analyzed the activity assays. LE and DRA interpreted the activity assays. All authors were involved in data analysis. HZ and JS wrote the manuscript with input from all authors.

## Data and materials availability

The cryo-EM maps and structure coordinates have been deposited in the Electron Microscopy Data Bank and Protein Data Bank under the accession codes EMD-75251 and PDB 10KO (Rab10–LRRK2 monomer); EMD-74979 and PDB 9ZZ1 (Rab10–LRRK2 dimer); EMD-75252 and PDB 10KQ (Rab10–LRRK2^RCKW^−DARPin E11 monomer); EMD-75256 and PDB 10KS (LRRK2^RCKW^−DARPin E11 dimer); EMD-75402 and PDB 10RF (LRRK2^RCKW^−DARPin E11 tetramer); EMD-74818 and PDB 9ZUJ (Rab8A–LRRK2 dimer); EMD-75007 and PDB 9ZZL (Rab43−LRRK2); EMD-74966 and PDB 9ZYY (Rab5A–LRRK2); EMD-75455 and PDB 10TF (GABARAP−LRRK2). Plasmids encoding constructs used for cryo-EM in this study are available upon request.

All the primary immunoblotting data presented in this study have been deposited in the Zenodo data repository (10.5281/zenodo.7601062). Plasmids were generated inhouse at the MRC Protein Phosphorylation and Ubiquitylation Unit at the University of Dundee and can be requested through the MRC-PPU reagents website (https://mrcppureagents.dundee.ac.uk/).

